# NMDARs in Granule Cells contribute to parallel fiber - Purkinje cell synaptic plasticity and motor learning

**DOI:** 10.1101/2021.01.12.426386

**Authors:** Martijn Schonewille, Allison E. Girasole, Philippe Rostaing, Caroline Mailhes-Hamon, Annick Ayon, Alexandra B. Nelson, Antoine Triller, Mariano Casado, Chris I. De Zeeuw, Guy Bouvier

**Affiliations:** Department of Neuroscience, Erasmus MC, Rotterdam, The Netherlands; Neuroscience Graduate Program, UCSF, San Francisco, CA 94158, USA; Department of Neurology, UCSF, San Francisco, CA 94158, USA; Kavli Institute for Fundamental Neuroscience, UCSF, San Francisco, CA 94158, USA; Weill Institute for Neurosciences, UCSF, San Francisco, CA 94158, USA; Department of Neurobiology, Harvard Medical School, Boston, MA, USA; Howard Hughes Medical Institute, Harvard Medical School, Boston, MA, USA; Cell Biology of the Synapse. Ecole Normale Supérieure, IBENS, Paris, F-75005, France. CNRS, UMR 8197, Paris, F-75005, France. INSERM, U 1024, Paris, F-75005, France. PSL, Paris, France; Ecole Normale Supérieure, IBENS, Paris, F-75005, France. CNRS, UMR 8197, Paris, F-75005, France. INSERM, U 1024, Paris, F-75005, France. PSL, Paris, France; Netherlands Institute for Neuroscience, Royal Dutch Academy of Arts &Sciences, Amsterdam, The Netherlands; Department of Physiology, University of California, San Francisco, San Francisco, CA, USA; Howard Hughes Medical Institute, University of California, San Francisco, San Francisco, CA, USA

## Abstract

Long-term synaptic plasticity is believed to be the cellular substrate of learning and memory. Synaptic plasticity rules are defined by the specific complement of receptors at the synapse and the associated downstream signaling mechanisms. In young rodents, at the cerebellar synapse between granule cells (GC) and Purkinje cells (PC), bidirectional plasticity is shaped by the balance between transcellular nitric oxide (NO) driven by presynaptic NMDA receptor (NMDAR) activation and postsynaptic calcium dynamics. However, the role and the location of NMDAR activation in these pathways is still debated in mature animals. Here, we show in adult rodents that NMDARs are present and functional in presynaptic terminals where their activation triggers nitric oxide signaling. In addition, we find that selective genetic deletion of presynaptic, but not postsynaptic, NMDARs prevents synaptic plasticity at parallel fiber-Purkinje cell (PF-PC) synapses. Consistent with this finding, the selective deletion of GCs NMDARs affects adaptation of the vestibulo-ocular reflex. Thus, NMDARs presynaptic to PCs are required for bidirectional synaptic plasticity and cerebellar motor learning.

**Significance Statement:** Learning depends on synaptic plasticity. The signaling mechanisms that control induction of plasticity determine the learning rules at the specific synapse involved. Moreover, the relationship between the activity patterns of synaptic inputs and the type, direction, and level of plasticity induced may evolve during development. Here, we establish a key link between NMDA receptor activation presynaptic to cerebellar Purkinje cells, downstream signaling mechanisms, and the ability of adult animals to learn a cerebellar motor task.

## Introduction

The ability of an organism to adjust its behavior to environmental demands depends on its capacity to learn and execute coordinated movements. The cerebellum plays a central role in this process by optimizing motor programs through trial-and-error learning (1). Within the cerebellum, the synaptic output from granule cells (GCs) to Purkinje cells (PCs) shapes computational operations during basal motor function and serves as a substrate for motor learning (2). Several forms of motor learning depend on changes in the strength of the parallel fiber (PF), the axon of GCs, to the PC synapse (3, 4).

In the mammalian forebrain, synaptic plasticity typically relies on postsynaptic NMDAR activation, which alters AMPA receptor (AMPAR) turnover at the postsynaptic site (5). However, this may not extend to the cerebellar synapse between GCs and PCs, since no functional postsynaptic NMDARs have been identified in young or adult rodents (6, 7). Pharmacological approaches, however, have shown that both long-term depression (LTD) and long-term potentiation (LTP) induction depend on NMDAR activation at the PF-PC synapse in young rodents (8–12). Hence, the alternative mechanisms for NMDAR-dependent synaptic modulation may involve presynaptic NMDARs activation (12-15; for review: 16, 17). Indeed, cell-specific deletion of NMDARs in GCs abolished LTP in young rodents (12). In addition to NMDARs, PF-PC synaptic plasticity also requires nitric-oxide (NO) signaling (19–21). As nitric-oxide synthase (NOS) is expressed in GCs, but not in PCs (22), the activation of presynaptic NMDARs might allow Ca^2+^ influx that activates NO synthesis, which in turn may act upon the PCs. However, in the mature cerebellum, the existence of presynaptic NMDARs on PFs and the role of NO in PF-PC plasticity remains a matter of debate. Previously, we have proposed that the activation of putatively presynaptic NMDARs in young rodents is necessary for inducing PF-PC synaptic plasticity without affecting transmitter release (8, 9, 11, 12). More recently, it has been shown that a subset of PFs express presynaptic NMDARs containing GluN2A subunits and that these receptors are functional (11, 12). Thus, in contrast to their role at other synapses, at least in young rodent, presynaptic NMDARs as part of the PF-PC synapses might act via the production of NO to induce postsynaptic plasticity, without altering neurotransmitter release (9, 11, 12, 19–23). However, a causal link between NMDARs activation in PFs, NO synthesis, and synaptic plasticity induction is still missing.

In the cerebral cortex, the expression of presynaptic NMDARs is developmentally regulated (24, 25). However, little is known about the presence and function of presynaptic NMDARs in adult tissue. In the adult cerebellum, PCs only express postsynaptic NMDARs at their synapse with climbing fibers (CFs) (18). It has been proposed that the activation of these receptors could have heterosynaptic effects during PF-PC long-term depression (LTD). This mechanism would explain why LTD in adults depends on NMDARs. According to this model, presynaptic NMDARs would be a transient feature of developing tissue and not necessary for induction of synaptic plasticity and motor learning in adult animals (18).

Here, we combine electron microscopy, 2-photon calcium imaging, synaptic plasticity experiments, and behavioral measurements, to show that presynaptic NMDARs are not developmentally regulated but are required for cerebellar motor learning in adults. We demonstrate that presynaptic NMDARs are present and functional in PFs of mature rodents. By specifically deleting the NMDAR subunit GluN1 either in the post-(PC) or the pre-synaptic cells (GCs), we demonstrate that NMDAR activation in GCs plays a key role in bidirectional synaptic plasticity and in vestibulo-ocular reflex (VOR) adaptation, an important paradigm for testing cerebellar motor learning (26–28). In contrast, NMDARs in PCs are neither involved in PF-PC synaptic plasticity nor required for cerebellar motor learning.

## Results

### NMDARs are presynaptically expressed and are activated by high-frequency stimulation of parallel fibers

First, we established the presence of presynaptic NMDARs in PFs of adult rodents by performing both pre-embedding and post-embedding electron microscopy immunohistochemistry, using antibodies directed against either GluN1 or GluN2 subunits. Pre-embedding immunoperoxidase staining revealed numerous GluN2 (Fig. 1*A*) and GluN1 (Fig. 1*B*) reactive profiles in PF varicosities facing small postsynaptic elements. The latter had cytological characteristics of PC dendritic spines (29). Pre-embedding immunogold labelling of GluN2 (Fig. 1*C*) and GluN1 (Fig. 1*D*) and distance measurements (Fig. 1*E*) indicated that particles are predominantly observed at the edge of the presynaptic active zones (82.5% are out of the active zone with 50% of the total at less than 125 nm from its edge). The same result was observed with post-embedding techniques (Fig. 1*F*). Next, we tested whether PF varicosities showed activity-dependent NMDAR-associated calcium transients. To follow calcium dynamics in PFs, we injected adeno-associated viruses carrying floxed GCaMP6f and td-Tomato in the cerebellar vermis of α6-Cre mice; α6 is a promoter specific of cerebellar GCs (30). Three to four weeks post viral injection, td-Tomato fluorescence was observed to the GC somatodendritic compartment in the GC layer and their PFs axons in the molecular layer (Fig. 1*G*). We stimulated PFs in cerebellar transverse slices using an extracellular electrode and measured calcium transients 50 to 250 μm from the stimulated site. This distance ensured the absence of presynaptic calcium dynamics perturbations (12). We stimulated PFs in bursts every 15 sec (25 pulses at 200 Hz) to show the presence of functional NMDARs in adults PF boutons. PFs stimulation produced calcium transients of variable amplitude between varicosities. After a stable baseline was achieved, we blocked NMDARs by applying a mix of D-APV (150 μM) and buffered Zn^2+^ (300 nM). In the presence of these drugs, the mean calcium transient amplitude was reduced (Fig. 1*H-J*, 95.3 ± 1.1% of baseline, p = 7.5e-5, n = 107 detected putative boutons). Upon drug washout, calcium transient amplitudes returned to baseline values (98.2 ± 0.8% of baseline, p = 4e-3 vs NMDAR blockers; p = 0.02 vs control). The calcium transient amplitude presented a skewed distribution only in the presence of the NMDAR blocker D-APV (Fig. 1*J*), consistent with the heterogeneous expression of NMDARs at these varicosities reported in young animals (11, 12). Thus, in mature PF boutons, presynaptic NMDARs are located at the periphery of the active zone and may serve a functional role.

**Figure 1:**
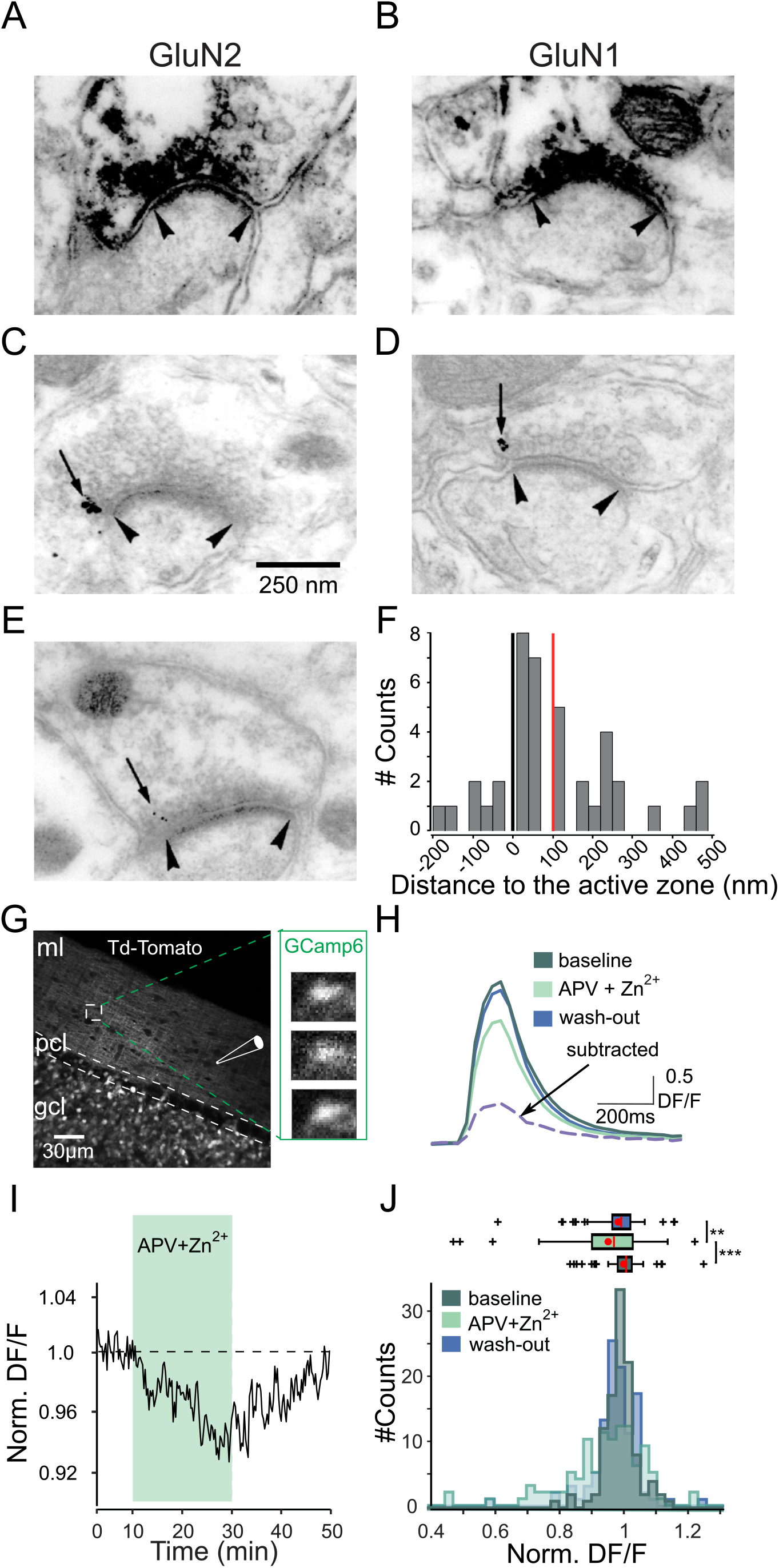
NMDARs are present and functional in PF varicosities. **(A-F)** Electron immunohistochemistry reveals the presence of GluN1 and GluN2 on presynaptic sites of PF-PC synapses. NMDAR labelling of profiles presynaptic to dendritic spines with antibodies recognizing GluN2 (A, C, E) or GluN1 (B, D) subunits. GluN2 (A) or GluN1 (B) subunit immunoperoxidase deposit is detected in the presynaptic element of asymmetric synapses. The presynaptic particles associated with GluN2 (C) or GluN1 (D) antigenic determinants (arrows) are detected at the edge (arrowheads) of the active zone following pre-embedding immunogold labelling. With post-embedding immunogold labeling GluN2. **(E)** associated particles are also found at the edge of presynaptic actives zones showing that the access to antigenic determinant was not limited by the cytoskeleton of the presynaptic differentiation. **(F)** Gold particle quantification of the distance to the edge of the active zone for GluN2: values along the x-axis represent the distance between the edge of the synaptic complex and the nearest immunogold labeling (mean = 100.1 nm (red), median = 50 nm, SEM = 24.3, n = 41), negative values represent particles within the active zone. Note: 82.5% of the gold particles are outside the active zone (right to the dark vertical line). **(G-J)** Calcium imaging of GCaMP6f-expressing PF varicosities. **(G)** Acousto-optic deflectors based 2-photon snapshot projection (1 plane, 10 images) of a granule cell expressing td-Tomato. PFs are directly stimulated in the molecular layer (ml) and images are recorded at least at 50 μm from the stimulation point (gcl: granule cell layer, pcl: Purkinje cell layer, ml: molecular layer). Enlarged: varicosity calcium image example before (top), during (middle), and after (bottom) blocking NMDARs (top to bottom, respectively). **(H)** Calcium transient example of another labeled varicosity (25-30 PFs stimulations at 200Hz). Baseline, APV and Zn^2+^ application, wash-out, and ([baseline]-[NMDAR blockade]) subtraction (purple dashed). Note the NMDAR-blockade effect on the calcium transient. **(I)** Time course of the normalized ΔF/F in control conditions (n = 107 varicosities, 31 slices from N = 14 mice). 150 μM D-APV and 300 nM Zn^2+^ were bath applied from minute 8 to 28. **(J)** Normalized ΔF/F data histogram. Box plot of normalized data comparing the signal in control conditions, during NMDAR block, and after washout (blue). Mean (red dots) and median (red lines) are shown. Statistical significance was tested using Wilcoxon test (*** p < 0.001, ** p < 0.01).

### Presynaptic, but not postsynaptic, NMDARs are required to induce PF-PC plasticity

In young rodents, we have previously shown that PF-PC synaptic plasticity required high frequency burst activation of PFs to recruit presynaptic NMDARs (11, 12). Moreover, we have previously confirmed that high frequency bursts also induce LTP in adult mice at the age of 2-3 months (31). Therefore, using bursts of 5 stimulations at 200 Hz every second (300 repetitions) of PFs, we first induced a postsynaptically-expressed potentiation of 227±19% compared to baseline (p = 4.4e-05; PPR = 98.8 ± 2.8% of baseline, p = 0.23; Fig. S1*A-D*). In contrast, bursts of 5 stimulations at 16.7 Hz every second (300 repetitions) of PFs did not result in a robust potentiation of the EPSC (125±5% of baseline; p = 3e-04 vs. 200 Hz condition, p = 0.09 vs. baseline, Fig. S1*A-D*) as shown in young rodents for both LTD and LTP (11, 12; respectively). The LTP induction at 200 Hz required NMDAR activation, as potentiation was blocked in the presence of D-APV (111 ± 7% of baseline; P = 4e-04 vs. control, P = 0.12 vs. baseline) (Fig. S1*A-D*). To determine the location of the NMDARs involved in LTP induction, we generated mice lacking NMDARs specifically in GCs by crossing α6-Cre with GluN1-floxed animals (GC-GluN1 ko, Fig. 2*A*, see Methods). Mice homozygous for the deleted GluN1 allele were viable and their cerebellum appeared to develop normally. High-frequency burst stimulation of the PFs (5 stimulations at 200 Hz every second, 300 repetitions) revealed that PF-PC LTP was impaired in these animals (109 ± 7 % vs 200 ± 40%, in slices from control animals GluN1 flox/flox without Cre, GC-GluN1 wild-type (wt), p = 3e-5; Fig. 2*B, D-F*). Likewise, PF-PC LTD was also abolished in these animals (110 ± 3% versus 70 ± 0.5 % in slices from control animals GC-GluN1 wt, p = 0.0009; Fig. 2*C-F*) using bursts of 2 stimulations at 200 Hz every second (300 repetitions) of PFs paired with high-frequency CF burst to induce LTD (31, see Methods). These results demonstrate that NMDARs expressed by GCs in adult animals are necessary for both LTP and LTD induction at PF-PC synapses. To determine whether NMDARs expressed by PCs also contribute to synaptic plasticity in mature mice, we produced mice lacking NMDARs specifically in PCs by crossing L7-Cre with GluN1 floxed mice (PC-GluN1 ko, Fig. 2*A*). L7 is a promoter that is specific for cerebellar PCs (32, see Methods). PF-PC LTP was indistinguishable from controls (206 ± 17 % versus 209 ± 26% in slices from control animals PC-GluN1 wt, p = 0.56; Fig. 2*B, G-I*) using high-frequency burst stimulation of the PFs (5 stimulations at 200 Hz every second, 300 repetitions). Likewise, PF-PC LTD was unaffected in slices from these animals (70 ± 4 % versus 76 ± 2 %, in slices from control animals GluN1 flox/flox without Cre, PC-GluN1 wt, p = 0.36; Fig. 2*C, G-I*) using bursts of 2 stimulations at 200 Hz every second (300 repetitions) of PFs paired with high-frequency climbing fiber burst to induce LTD (31, see Methods). Thus, the NMDARs necessary for LTP and LTD induction at mature PF-PC synapses are located presynaptically in the GCs and not in the postsynaptic PCs.

**Figure 2:**
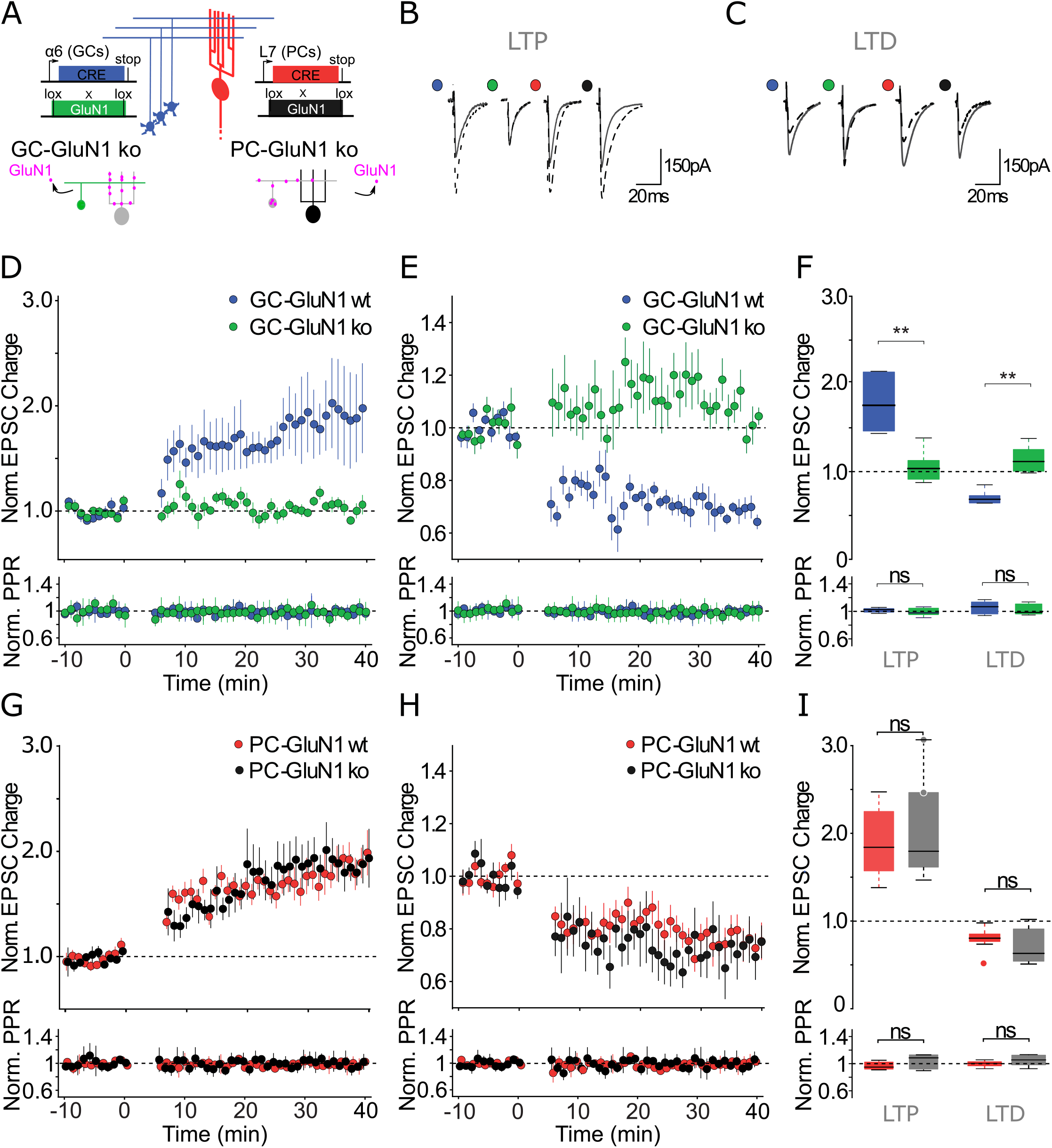
Synaptic plasticity in animals lacking NMDARs in specific neuronal populations. **(A)** Schematic representation of the strategy used to knock out the GluN1 gene either in cerebellar granule cells (GC-GluN1, wt:blue/ko:green) or in Purkinje cells (PC-GluN1, wt:red; ko:black) **(B, C)** Representative recordings before (grey) and after (dashed line) LTP **(B)** and LTD **(C)** induction. The colored dots beside the traces are the same as in (D-I). **(D, E)** Time course of the normalized EPSC charge (top) and PPR (bottom), in GC-GluN1wt (blue) and GC-GluN1ko (green) for LTP (D) and for LTD (E). **(F)** Normalized EPSC charge (top) and PPR (bottom) after plasticity induction (t = 30-35 min) for all individual experiments in GC-GluN1wt (LTP, n = 8 cells; LTD, n = 7 cells) and GC-GluN1ko (LTP, n = 10 cells; LTD, n = 6 cells). **(G, H)** Time course of the normalized EPSC charge in PC-GluN1 wt (red) and PC-GluN1 ko (black) for LTP (G) and for LTD (H). **(I)** Normalized EPSC charge (top) and PPR (bottom), after plasticity induction (t = 30-35 min) for all individual experiments in PC-GluN1wt (LTP, n = 10 cells; LTD, n = 7 cells) and PC-GluN1ko (LTP, n = 9 cells; LTD, n = 6 cells). PPR was not changed after synaptic plasticity induction (GC-GluN1: D-F, bottom; PC-GluN1: G-I, bottom). Boxes represent median (black), the upper and lower quartile of the distribution. To induce LTP, we use 5 stimulations at 200 Hz every second, 300 repetitions, while using bursts of 2 stimulations at 200 Hz every second (300 repetitions) of PFs paired with high frequency climbing fiber burst to induce LTD (see Methods). Statistical significance was tested using Wilcoxon test (ns: p > 0.05; ** p < 0.001).

In contradiction to our results, Piochon and colleagues (18) observed no effect of NMDAR blocking on LTP induction. Therefore, we tried to identify the experimental factors responsible for this difference. In the direct vicinity of the stimulation electrode (up to a few tens of microns), presynaptic (and presumably post-synaptic) calcium dynamics can be disturbed (12). Since the stimulation electrode is usually in the immediate vicinity of the recorded synapse in sagittal sections, the requirements for presynaptic NMDAR activation may be bypassed under these conditions. To check for a potential bias created by slice orientation, we tested plasticity induction in sagittal slices. Indeed, using the same conditions as Piochon and colleagues (18) (7 PF stimulations at 100 Hz, 300 repetitions at 1 Hz) and our LTP induction protocol used in horizontal slices (5 PF stimulations at 200 Hz every second, 300 repetitions), we successfully induced LTP (142 ± 11% of baseline; p = 0.0002; Fig. S1*E-G*; and 149 ± 10% of baseline; p = 6e-4; Fig. S1*E, G*; respectively). In contrast with horizontal slice orientation experiments, this form of LTP was not abolished by APV application (148 ± 16% of baseline, p = 0.003, p = 0.95 vs. sagittal control condition, Fig. S1*E-G*; and 139 ± 14% of baseline, p = 0.008, p = 0.43 vs. sagittal control condition, Fig. S1*E, G*; respectively). Therefore, we propose that the absence of an effect of NMDAR blockade reported by Piochon and colleagues (18) might be due to direct perturbation of the presynaptic calcium dynamic in PFs terminals.

To connect the requirement for high frequency stimulation of PFs to induce PF-PC plasticity and the kinetic properties of NMDARs, we investigated the involvement of presynaptic NMDARs containing the GluN2A subunits in PF-PC synaptic plasticity. It has been demonstrated that presynaptic NMDARs at PF-PC synapses in young animals show pharmacological properties characteristic of GluN2A-containing receptors (11, 12, 33), which may largely determine receptor kinetics (24, 25, 34). To determine if NMDARs containing GluN2A subunits are involved in LTP induction in adult mice, we investigated LTP induction in the global GluN2A-KO mice (35). In these mutants lacking GluN2A subunits, LTP could not be induced with our standard protocol: 5 PF stimulations at 200 Hz, 300 repetitions (106 ± 15% of baseline, p = 0.21, Fig. S1*B, D*), in contrast to their wild-type littermates (177 ± 18% of baseline). This result shows that GluN2A-containing NMDARs are required for plasticity induction at PF-PC synapses in adults. These data are consistent with earlier demonstrations in younger mice (12), highlighting persistent expression patterns of NMDAR subunits at PF-PC synapses during postnatal development and beyond. Thus, PF-PC bidirectional synaptic plasticity in the mature cerebellum is driven by NMDARs containing GluN2A subunits.

### Nitric oxide dependent LTP requires activation of GCs expressing NMDARs

Both LTP and LTD induction depend on nitric oxide (NO) signaling in young rodents in the cerebellum (19–21, 36). However, this finding has yet to be shown in adult animals. To test the involvement of NO signaling in adult LTP, we applied our standard LTP induction protocols in the presence of L-NAME (100 μM), a specific antagonist of NOS. Under these conditions LTP was abolished (101 ± 5% of baseline, p = 0.47; Fig. S2*A-C*). NO synthesis during PF bursting activity may result from activation of presynaptic NMDARs (9, 11, 12) and/or from activation of receptors located in interneurons (10, 37). To understand the link between NMDAR activation and NO synthesis and determine the cell types involved, we used an electrochemical probe to measure NO production during high-frequency burst stimulation in the molecular layer (see Methods). We stimulated PFs with a protocol identical to the previous configuration used to image NMDAR-dependent calcium signals (Fig. 1). This protocol resulted in a significant enhancement of the current recorded at the electrode (Fig. S2*D, E*; 112.8 ± 32.8pA, p = 8e-5, GC-GluN1 wt). The signal was specific for NO production, as it was absent in the presence of L-NAME (Fig. S2*D, E*; −33.5 ± 34.1pA, p = 0.18). Moreover, NO production was due to NMDAR activation, because it was also abolished in the presence of APV (Fig. S2*D, E*; −68.1 ± 27.1pA, p = 1.3e-4 APV vs control). Finally, the signal was absent in slices from mice lacking GluN1 in GCs (Fig. S2*D, E*; −10.4 ± 27.2pA, n =14, wt vs. ko, p = 0.003). Therefore, we conclude that high-frequency activity of PFs results in activation of their NMDARs and that subsequent NO production participates in the induction of plasticity within the molecular layer.

### Cerebellum-dependent basal motor behavior is virtually unaffected after ablation of NMDARs from GCs or PCs

To test the specific contributions of NMDARs in GCs and PCs in basal reflexive eye movements, we subjected animals lacking NMDARs specifically in either GCs or PCs to a cerebellum-dependent behavioral assay involving compensatory eye movements. Vestibular and full-field visual inputs are known to drive compensatory eye movements via the vestibulo-ocular reflex (VOR) and optokinetic reflex (OKR), respectively (38, 39). While the VOR compensates for head movements with contra-directional eye movements, the OKR causes the eyes to follow a moving visual field while the head is stationary. These reflexes work in conjunction to generate the visually-enhanced VOR (or VVOR), which allows for the maintenance of a stable image on the retina while an animal moves through its environment. To preserve optimal stabilization throughout life the VOR is subject to adaptation based on visual feedback, a process that depends on the cerebellar cortex (3). Therefore, we used these different behavioral paradigms to evaluate the contribution of pre- and postsynaptic GluN1 receptors to motor performance. We tested the baseline OKR, VOR, and VVOR in wt, PC-GluN1 ko mice, and GC-GluN1 ko mice (Fig. S3*A*). In PC-GluN1 ko mice, eye movements evoked by sinusoidal rotation of the visual field (OKR) or table (VOR) did not differ in gain (reflecting amplitude) or phase (timing) from those of controls (Fig. S3*B*, OKR and VOR: all p > 0.5). In the VVOR, combining both inputs, the gain was also not affected (p = 0.55), but there was a small difference in phase (p = 0.010, Δphase = 1.1 ± 0.1° across tested frequency range). Likewise, ablating GluN1 from GCs mildly affected compensatory eye movements (Fig. S3*C*). VOR phase was lower in GC-GluN1 ko mice (p = 0.043, Δphase = 6.2 ± 2.2°), whereas all other parameters did not differ from those of controls (all p > 0.3), despite a trend in the gain of the VVOR to be lower in GC-GluN1 ko mice (p = 0.056). Taken together, ablating GluN1 from PCs or GCs has minimal effects on baseline performance of the VOR and OKR, suggesting that NMDARs and NMDAR-dependent plasticity are only marginally important for basal cerebellum-related motor behavior.

### VOR adaptation deficits are more pronounced after ablating NMDARs from GCs

Genetic interventions, particularly those targeted at components of plasticity processes, typically affect motor learning more than motor performance (28, 40, 41). To investigate the role of GCs and PCs NMDARs in cerebellar motor learning, we first tested the contribution of GluN1 to short-term learning by subjecting mice to a VOR learning paradigm designed to decrease VOR gain. Five sessions of sinusoidal rotation of the mice and the visual input in the same direction at the same amplitude (inphase) for 10 min resulted in a progressive decrease of VOR gain in control mice (recorded in dark, p < 0.001, before vs. after 50 min; Fig. 3*A, B*). This type of short-term cerebellar learning was not affected in PC- or GC-GluN1 ko mice (both p > 0.14 vs respective controls; Fig. 3*A, B*). Next, we tested mice on a longer-term learning paradigm by continued training the following day (after mice remain in the dark for 23 hours) with the same parameters. However, the amplitude of visual stimulus was increased. This paradigm was designed to reverse the direction of the original VOR (i.e. VOR phase adapt towards 180°). Over three days of reversal training, VOR phase increased from values ≤ 30° to an average maximum of 144 ± 15° in control and 132 ± 13° in PC-GluN1 ko mice (Fig. 3*A*). This phase increase was comparable between groups (day 2, 3, and 4; all p > 0.3). While the VOR gain increased again after the phase had reversed, this gain increase was significantly less pronounced in PC-GluN1 ko mice (day 4, p = 0.011), indicating a subtle impairment in VOR learning (Fig. 3*A*). In contrast, the effects in GC-GluN1 ko mice were more severe. These mice did not achieve the same phase-shift as control mice, reaching an average maximum of 58 ± 10° (GC-GluN1 ko) compared to 115 ± 16° (wt; Fig. 3*B*). This difference was significant throughout the training period (p < 0.015 for days 2, 3, and 4). In addition to its role in VOR adaptation, the cerebellum is also important for adaptation of the OKR. Prolonged exposure to sinusoidally rotating visual input, combined with or without a conflicting vestibular input, will result in an adaptive increase of OKR gain (40). OKR gains after the reversal training were significantly higher in all mice, but this increase was attenuated in GC-GluN1 ko mice (Fig. 3*C*, p = 0.045), whereas there was no impairment in PC-GluN1 ko mice (p = 0.27). The presence of an evident phenotype in both cerebellum-dependent learning paradigms, VOR phase-reversal and OKR gain-increase, confirms the functional relevance of NMDARs in GCs in cerebellum-dependent learning.

**Figure 3:**
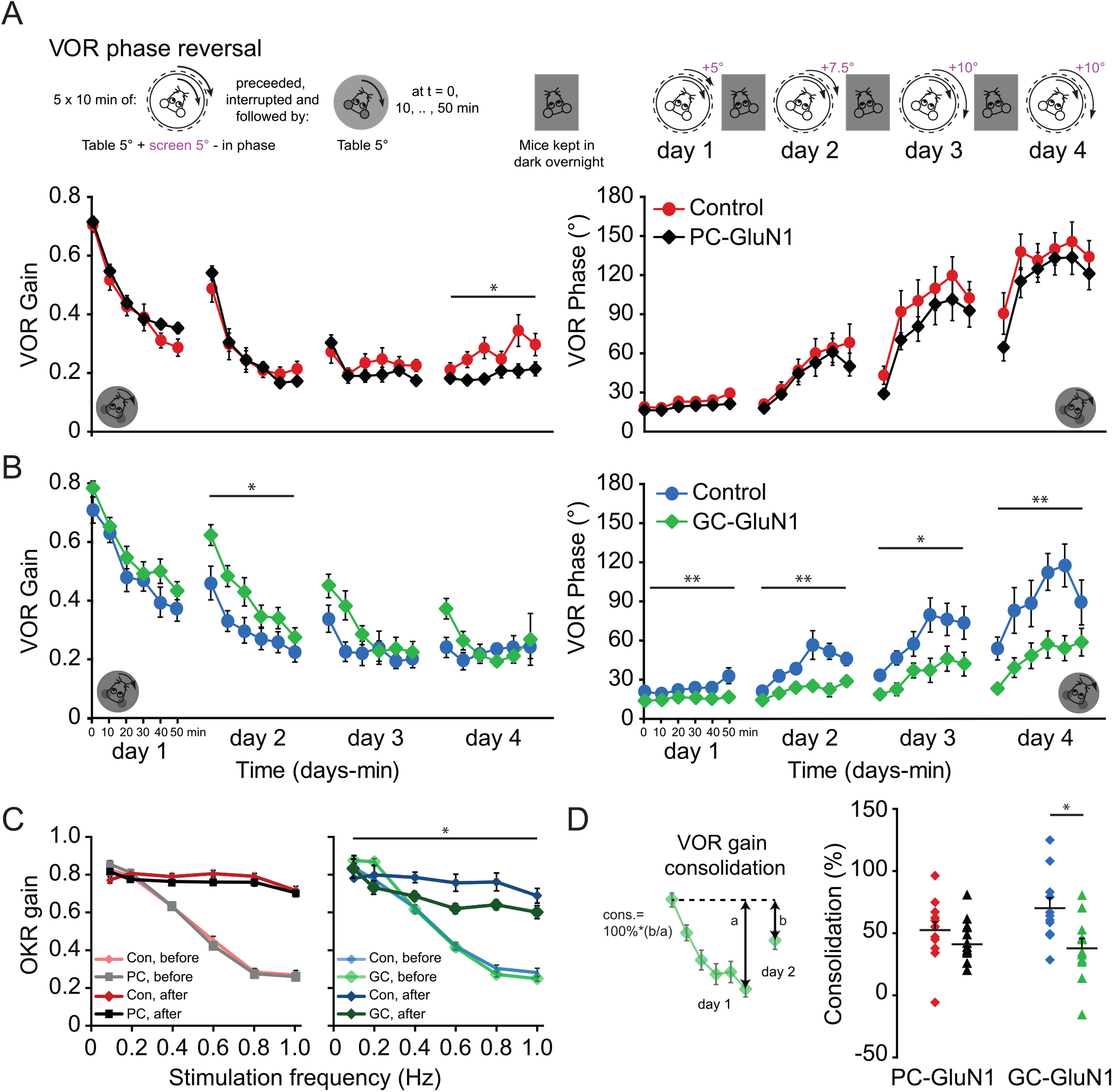
Cerebellum-dependent VOR phase reversal is affected in mice lacking NMDARs in GCs. **(A)** Top panels: Schematic representation of the VOR phase reversal experiment. Mice were subjected to a mismatched combination of vestibular and visual input. VOR phase reversal is induced by five 10 min training sessions during which the visual stimulus is rotated in-phase with the vestibular input (amplitude, 5°) at increasing amplitudes (day 1, 5°; day 2, 7.5°; day 3-4, 10°). Bottom panels: VOR phase reversal training typically starting with a decrease of VOR gain (day 1), followed by the increase of the VOR phase (aimed at a phase value of 180°, day 2-4) and, when VOR phase is sufficiently reversed, the increase in the VOR gain (day 4). PC-GluN1 ko mice show VOR phase reversal with similar phase values as controls (both N = 12 mice), but are impaired in the VOR gain increase after reversal (day 4, p = 0.011). **(B)** Mice lacking NMDARs from GCs have a more pronounced deficit in that the phase increases more slowly (day 2, 3, 4: p = 0.001, 0.013, 0.006, respectively). Note that control littermates of the GC-GluN1 mutant mice appear to adapt their VOR slower than littermates of the PC-GluN1 mutant mice, presumably due to subtle batch and inter-experimental differences. **(C)** VOR phase reversal training also results in an enhanced gain of the OKR. This cerebellum-dependent adaptive change was also impaired in mice lacking NMDARs (N = 11 mice) in GCs (vs. 7 control mice, p = 0.045), but not in PC-GluN1 ko mice (both N = 11 mice, p = 0.27). **(D)** VOR gain consolidation, the percentage of learned response present on day 2 (b) relative to the adaptive change on day 1 (a) (see inset), was not affected in PC-GluN1 mice (p = 0.38, both N = 12 mice, unpaired Students *t*-test). In contrast, impaired long-term adaptation could be linked to a deficit in consolidation in GC-GluN1 ko mice, compared to controls (p = 0.011, both N = 11 mice, unpaired Students *t*-test). (D) Error bars denote SEM (please note that error bars can fall within symbols), * p < 0.05, ** p < 0.001.

Overnight consolidation, the stabilization of acute adaptive changes to long-term modifications, is crucial for persistent motor adaptation (42, 43). To uncover whether the GC-GluN1 phenotype is linked to the ability to adapt during training or the ability to store and maintain the adaptation overnight, we tested the consolidation of the adaptive changes (44). As this measure is relatively sensitive to noise, we compared the most stable, consistent change: i.e. the difference between the VOR gain-decrease during the first day and that on the second day. Consolidation, the percentage of change persisting overnight, was impaired in GC-GluN1 ko mice (Fig. 3*D*, 39 ± 8% vs. 71 ± 8% in controls, p = 0.011), implying that at least part of the deficit can be explained by impaired consolidation. Not surprisingly, consolidation was not impaired in PC-GluN1 ko mice (Fig. 3*D*, 41 ± 6% vs. 52 ± 7% in controls, p = 0.38). This selective impact on consolidation underscores the longterm impact of the GluN1-dependent mechanisms hypothesized above.

## Discussion

Here we present immunohistochemical and functional evidence for the presence of presynaptic NMDARs at PF-PC synapses in the cerebellum of adult rodents. As in young rodents, these receptors are required to induce both postsynaptic forms of synaptic plasticity, LTP and LTD. Using cell-specific deletion of NMDARs either in GCs or PCs, we demonstrate that only GC NMDARs are robustly involved in PF-PC synaptic plasticity and VOR adaptation. Counter to previous theories, our results support the idea that these receptors are persistently expressed in the adult cerebellum and that the roles of PC NMDARs in PF-PC synaptic plasticity and cerebellar learning should be reconsidered.

### Location of presynaptic NMDARs

Presynaptic receptors are often found within the active zone where they modulate synaptic transmission and plasticity. This phenomenon is seen in various types of presynaptic receptors throughout the brain (13–15), including GABA receptors (45), NMDARs (16, 17), and kainate receptors (46), all of which are found on PF terminals (47). However, presynaptic NMDARs at the cerebellar PF-PC synapse control plasticity in a distinct manner, these receptors participate in the induction of plasticity by promoting NO synthesis without changes in transmitter release (9, 11, 12, 37). This stands in marked contrast to other central synapses like in hippocampus (48, 49) or cerebral cortex (49, 50). How the activation of a calcium-permeable presynaptic receptors at the cerebellar PF-PC synapse results in calcium-dependent signaling (NO synthesis) without interfering with transmitter release has been a puzzle, leading to doubts regarding the existence of such receptors. Here, we propose an explanation for this paradox. Immunogold staining performed in adult rats shows that most of the presynaptic NMDARs are not located within the active zone, but at its periphery, thus capable of being active without affecting calcium dynamics at release sites. For comparison, this location is beyond the 30 nm coupling distance between calcium influx through voltage-dependent calcium channels and the release sensor (51). While our immunogold staining were obtained from adult rats, these results are not only consistent with our mice experiments using various other techniques, but also with several studies reporting an absence of release probability change after PF-PC synaptic plasticity induction in mice and rats (8–12, 18–21, 23, 31, 36–39, 57–59, 63–64). Therefore, it is likely to observe a similar NMDARs distribution at the periphery of PFs boutons in both rats and mice.

As demonstrated in young rodents, we show that PF-PC synaptic plasticity is NO-dependent. While NMDARs-dependent NO synthesis has been shown in young rodents, the origin of NO is still a matter of debate (9–12, 33, 37). Here, using specific deletion of NMDARs in PFs, we demonstrate that NMDAR activation leads to NO production, suggesting that the recruitment of indirect NMDAR-dependent signaling process from other cell-types is unlikely. This result suggests a tight coupling between NMDARs and NOS that may minimize crosstalk between the Ca^2+^ entry involved in transmitter release and the Ca^2+^signaling implicated in plasticity induction through the NMDAR-NOS system. These results are also consistent with the involvement of NO in PF-PC LTP described in this study, previous *in vitro* studies reporting a role of NO in LTD in adult rodents (52, 53), and visuo-motor behavior studies showing a learning impairment in NOS knockout mice (54).

### Role of presynaptic NMDARs in plasticity at the PF to PC synapse

We have previously described that presynaptic NMDARs are required to induce LTD (11) and LTP (12) at PF-PC synapses in cerebellar slices from young rodents. Our data demonstrates that both LTP and LTD induction are still NMDAR-dependent in adult animals. These results differ with previous reports suggesting a role of other cell types in PF-PC synaptic plasticity (18, 19). Using cell-specific deletions of the GluN1 subunit either in GCs (α6-Cre mice) or PCs (L7-Cre mice), we show that only the NMDARs expressed in GCs, and not those in PCs, are involved in both LTP and LTD at the PF-PC synapse. We propose two explanations for this discrepancy. First, in the parasagittal slice configuration used in other studies (18, 19), the proximity of the extracellular stimulation electrode to the presynaptic elements may perturb the calcium dynamics, thereby potentially bypassing the requirement for NMDAR activation on PFs (12). Consistently, we also found that NMDARs activation is not required to induce PF-PC LTP in sagittal cutting orientation (Fig. S1). Interestingly, using specific deletion of the GluN1 subunits in adult mice PCs, Kono and colleagues also reported that PF-PC LTD is independent of NMDARs expressed in PCs. Instead, they proposed that NMDARs located on molecular layers interneurons, and not on GCs, are required to induce PF-PC LTD (19). While the NMDAR subunits profile is not consistent with a role of molecular layer interneuron in PF-PC plasticity in young rodents (11, 12, 33), it remains to be determined whether molecular layer interneuron might also be involved in PF-PC synaptic plasticity using more physiological experimental conditions in adult rodent. The Second, climbing fibers (CFs) *in vivo* fire in bursts at a high frequency (reaching 400 Hz: 55, 56), which is well above the frequency of stimulation used by Piochon and colleagues. We induced LTD using a high frequency burst of the CF 100 ms after the PF burst, as described previously *in vitro* (31, 57) and *in vivo* (56, 58). This bursting parameter may be crucial, since calcium spike propagation in the PC dendritic arbor depends on membrane excitability (59). Therefore, a gentle hyperpolarization of the dendrite due to CF-NMDARs blockade could decrease the probability of propagation and consequently impair LTD induction (12, 60). This phenomenon is of potential physiological interest as it may attribute a subtle role in dendrite excitability to the NMDARs present at the CF-PC synapse. Finally, even though the experimental conditions such as age, recording temperature, slice orientation, and protocol for plasticity induction substantially differed among the various cerebellar studies, many of them reported a relatively low amplitude (~10-20 %) of PF-PC LTP (61, 38, 62). As this LTP is 5 to 10 times smaller than our LTP in control conditions, we could also imagine that a small fraction of our LTP does not depend on NMDARs. Taken together, presynaptic, but not postsynaptic, NMDARs are essential to induce bidirectional synaptic plasticity at the PF-PC synapse in adult mice.

### Motor performance and motor learning

The robust cellular effects on synaptic plasticity that we have described *in vitro* raise the question to what extent ablating NMDARs affects motor behavior. Here, we demonstrate that 1) ablating NMDARs from neither GCs nor PCs prominently affects motor performance, 2) in line with the absence of an effect on PF-PC synaptic plasticity, ablating NMDARs from GCs, but not from PCs, impairs VOR adaptation, and 3) the combined deletion of LTD and LTP attenuates, but does not completely block, cerebellar learning. By revealing these behavioral phenotypes in GC-specific versus PC-specific GluN-mutants, targeting two connected structures separately, our data expand upon those that have been obtained in global mouse mutants. Indeed, globally deleting GluN2A has also been found to attenuate, but not block, VOR phase-reversal learning (40), yet this study did not allow for any interpretation on the brain region or cell types involved in the learning process. Moreover, the additional value of using cell-specific mutants is also evidenced by interpreting the impact on motor performance. Whereas we were able to show in our GC- and PC-specific mutants that NMDARs have a minimal role in motor performance, earlier studies in global, double GluN2A/C knockouts were found to have severe motor performance deficits in the rotarod test (63), possibly resulting in misleading hypotheses. Using conditional mouse models in which GluN1 was deleted from parvalbumin (PV) expressing neurons, including PCs and molecular layer interneurons (MLIs), Kono and colleagues. (2019) observed deficits in OKR gain increase, that were absent in mice in which GluN1 was deleted from PCs only. It should be noted however, that deletion of GluN1 from PV-expressing neurons could also affect other circuits (64, 65, 66), including e.g. visual cortex that is known to also be required for OKR plasticity (67). In the same study, deletion of GluN1 from GCs did not cause a significant deficit in the single day of training (19), which is in line with the absence of a phenotype in GC-GluN1 mutant mice in the first day of training in our study. An extended training paradigm potentially could have revealed a deficit, as we found in our current study that long-term training is impaired. This deficit in long-term learning could in part be explain by a deficit in consolidation early in training, although the absence of a robust effect on subsequence days and relatively high consolidation in GC-GluN1 control mice compared to the PC-GluN1 control littermates, suggest that other factors contribute to the deficits as well.

Even though cell-specific mutants do not suffer from the limitations of the classical global mutants in that they allow for correlational interpretations at the level of brain region and cell type, they are equally limited when it comes to unraveling correlations at the subcellular level. As our cell-specific manipulation in the GC-specific mouse line ablates all NMDARs expressed in GCs, the effects on their ability to show cerebellar motor learning might be caused not only by the deficits at their axons (i.e., PF terminals), but partially by an absence of NMDARs at the level of their dendrites. Indeed, the connection between mossy fibers (MF) and GC dendrites is glutamatergic and may also undergo NMDAR-dependent LTP (68). Therefore, the deficits in cerebellar learning observed in our study in the α6Cre-GluN1 mice could in principle also be related to the putatively impaired induction of MF to GC LTP. The exact contribution of each of these two forms of synaptic plasticity (i.e., upstream, NMDA-dependent MF to GC plasticity and downstream PF to PC plasticity) to the motor learning deficits can only be determined with a technique that would allow for a specific deletion at the subcellular level, targeting dendrites and axons separately. Since such technology is currently not available to the best of our knowledge, we cannot reliably estimate the individual contributions of these two cellular processes to cerebellar learning behavior. Even so, we would like to highlight additional arguments why presynaptic NMDARs at the PF-PC synapse may at least contribute to cerebellar learning. First, the effects of GluN1 deletion in GCs, not only on LTD but also on LTP at the PF-PC synapse were dramatic. Second, the behavioral phenotypes observed in the mutants with a global deletion of GluN2 subunits (40, 68) may also result from a comparable effect on PF-PC synaptic plasticity, as we found that GluN2A is also required for PF-PC synaptic LTP (Fig. S1*B, D*). Third, VOR adaptation is probably mainly driven by increases in simple spike activity and LTP (69), which is congruent with the findings that VOR adaptation was affected to a similar extent by the selective loss of LTP (31, 70) and not affected by selective disruption of LTD alone (39, 71). Finally, whereas plasticity induction at PF to PC synapses by the presence and absence of CF inputs forms an ideal substrate for guided learning of specific inputs (66, 72), such level of input specificity controlled by teaching activity is lacking at the MF to GC synapse.

Taken together, we cannot rule out a potential contribution of MF-GC synaptic plasticity in VOR adaptation as evidenced by deficits observed after GluN1 deletion in GCs, but we can rule out an essential role of NMDARs expressed in PCs in this form of cerebellar motor learning. Moreover, the current study provides substantial supportive evidence for a contribution of NMDA-dependent PF to PC synaptic plasticity in VOR learning in adults, highlighting that GCs NMDARs may at least partially express NMDARs to mediate synaptic plasticity at their PF input to the PC dendrites, providing a diverse presynaptic machinery with ample means to fine-regulate the output of the cerebellum in the long-term (47, 72).

## Material and Methods

### Transgenic animals

Cellular specificity was verified by crossing L7Cre and BACα6Cre mice with floxed fluorescent reporter mice lines. See details in SI Material and Methods.

### Electron immunohistochemistry

Pre-embedding immunoperoxidase or immunogold and post-embedding immunogold detections were performed with specific anti-GluN1 mouse monoclonal antibody and anti-GluN2A rabbit polyclonal antibody. See details in SI Material and Methods.

### Electrophysiology in slices

Whole-cell patch-clamp recording in cerebellar slices from C57Bl6 mice (8-to 20 weeks-old) were performed the same as in ref. 31. Details are provided in SI Material and Methods.

### NO detection

NO efflux from cerebellar slices was monitored using a NO-selective amperometric microprobe with a 7 μm tip (WPI), similar to previous work (37). A constant voltage of 0.9V was applied using an Axon Multiclamp 700B amplifier. The probes were routinely cleaned in 0.1M H_2_SO_4_ to remove debris. The microprobe was positioned at least 30 μm away from the stimulating electrode to avoid spurious Ca-NO signaling (12). Before use, probes were equilibrated for 30 min in the incubation medium.

### Viral injection

2 to 3-month-old C57BL/6 Bac6-cre mice, in which recombinase is expressed exclusively in the cerebellar granule cells, were used. A mix of flexed GCaMP6f and td-Tomato adeno-associated viruses were stereotaxically injected into the cerebellar cortex. See details in SI Material and Methods.

### Calcium imaging

All experiments were performed using a custom-built random-access two-photon laser-scanning microscope. Details are provided in SI Material and Methods.

### Compensatory eye movement adaptation

At least 5 days after preparatory surgery, adult male mutant and control mice were head-fixed in the middle of a turntable with surrounding screen. Compensatory eye movements were evoked with rotational visual and/or vestibular input, and cerebellar motor learning was tested by mismatching these two inputs. Video-recorded eye movements were calibrated and gain and phase, representing the amplitude and timing of movement respectively, were determined. Details are provided in SI Material and Methods.

## Acknowledgements

This work was supported by the program “Investissements d’Avenir” from the French Government, implemented by Agence Nationale de la Recherche, references: ANR-10-LABX-54 MEMOLIFE, ANR-11-IDEX-0001-02 PSL* Research University. MS was funded by the Funding Dutch Scientific Organization – Life sciences (NWO ALW Veni) and ERC-Stg. CIDZ was supported by FP7-C7 European Commission, ZonMw, Netherlands Organization for Scientific Research (ENW-Klein), European Research Council (Advanced Grant, and Proof of Concept Grant), Medical NeuroDelta Programme, Topsector Life Sciences & Health (Innovative Neurotechnology for Society or INTENSE), and the Albinism Vriendenfonds Netherlands Institute for Neuroscience. GB was funded by Région Ile de France, FRM, and Labex MEMOLIFE. AEG was supported by the NSF GRFP and the UCSF Discovery Fellows Program. The funders had no role in study design, data collection and analysis, decision to publish, or preparation of the manuscript. We thank R de Avila Freire and L Post for technical assistance, and B Barbour, RS Larsen, PL Reeson, E Jones, M. Mukundan, and C Wang for comments on the manuscript.

## Notes

### Competing Interest Statement

The authors have declared no competing interest.

