## supplementary information for "NMDARs in Granule Cells contribute to parallel fiber - Purkinje cell synaptic plasticity and motor learning"

### Supporting Information

### SI Material and Methods

**Transgenic mice.** Cellular specificity was verified by crossing L7Cre and BACα6Cre mice with floxed fluorescent reporter mice lines, YFP and tdTomato, respectively. 2-3 month old mice were anesthetized with pentobarbital (60mg/kg) and intracardially perfused with 4% PFA in PBS. After O.C.T inclusion, brains were cut with a cryostat. YFP or tdTomato expressions were evaluated by immunohistochemistry combined with calbindin counterstaining of PCs, and DAPI for cell counting. L7-Cre x tdTomato floxed mice showed staining of 100±0% of PCs, 4.3±1.2% of molecular layer interneurons (MLI), and 2.3±1.3% of small neurons in the granule cell layer, presumably GCs (total number of cells sampled: 186 PCs, 1284 MLIs, and more than 6409 GCs; N=3 mice). BACα6Cre x YFP flox mice showed staining of the vast majority of granule cells. 4.8±1.4% of PCs also showed to be positive. No staining of MLI was observed (total number of cells sampled: 112 PCs and 840 MLIs; N=2 mice). The physiology experiments were performed blinded for the first 4 to 5 animals only, then for reasons of matching the number of animals the experimenter was no longer blind to the genotype. The behavioral experiments were entirely done blind to the genotype.

**Electron immunohistochemistry.** Adult Sprague Dawley rats (2-3 months old) were anesthetized with pentobarbital (60mg/kg) and intracardially perfused with 4 % paraformaldehyde (PFA) and 0.1% glutaraldehyde in phosphate buffered saline. After dissection, cerebella were kept overnight at 4°C in 4% PFA. For pre-embedding, frontal vibratome sections (100 μm) were cryoprotected and permeabilized by freezing and thawing and then incubated 24 hours at 4°C in anti-GluN1 mouse monoclonal antibody and anti-GluN2A rabbit polyclonal antibody (Euromedex, 1:1000 and 1:100 respectively). Immunoperoxydasic detection was performed as already described (Colin et al., 1998). For immunogold detection, vibratome sections were incubated in goat anti-mouse or goat anti-rabbit (both Nanoprobes, 1:100) nanogold coupled antibodies respectively. Gold particles were then stabilized, amplified and gold-toned. Sections were then dehydrated in graded ethanol and flat embedded in epoxy resin (Araldite, Ernest Fullam). For post-embedding immunogold labeling, 200 μm vibratome sections of cerebellum (same fixation as above) were rapidly frozen in liquid propane cooled in liquid nitrogen and freeze-substituted in methanol with 2 % uranyl acetate at –90°C for 30 hours. After raising the temperature from –90°C to –45°C, sections were rinsed in anhydrous methanol, infiltrated with Lowicryl HM20 resin (Fluka; Buchs Switzerland) and polymerized with UV light 48 hours at –45°C before increasing to room temperature. Ultrathin sections collected on nickel grids coated with formwar were etched and incubated with primary antibodies (anti-GluN1 1: 1000 and anti-GluN2A 1: 100). The secondary antibody used was a goat anti-rabbit IgG coupled to 10 nm colloidal gold particles (Biocell, 1:500, 1 hour at room temperature). Observations of ultrathin sections were performed with a JEOL 100 CX II electron microscope. Two rats were used for pre-embedding and two other rats were used for post-embedding labeling. Staining in every panel is present in elements having synaptic vesicles, a clear-cut signature of presynaptic buttons. These presynaptic elements make asymmetric contacts with postsynaptic elements having a postsynaptic density. The asymmetric distribution of the synaptic electrodense material is a signature of excitatory synapses. Furthermore, endoplasmic reticulum is often observed in the postsynaptic elements, indicative of Purkinje cell spines. Note that the quantification we present in Figure 1 has been performed on immunogold staining after pre-embedding in which we measure the tangential distribution of the gold particles.

**Electrophysiology in slices.** C57Bl6 mice (8- to 20 weeks-old) were anesthetized with isoflurane (Nicholas Piramal India Ltd.) and killed by decapitation. The cerebellum was rapidly dissected into cold solution containing (in mM): 230 sucrose, 26 $NaHCO_3$, 3 KCl, 0.8$CaCl_2$, 8$MgCl_2$, 1.25 $NaH_2PO_4$, 25 glucose supplemented with 50 μM D-APV to protect the tissue during slicing. Horizontal cerebellar acute slices (300 μM) were cut in this solution using a Campden Instruments 7000smz and stored at 32°C in a standard extracellular saline solution containing: 125 NaCl, 2.5

KCl, 2 $CaCl_2$, 1 $MgCl_2$, 1.25 $NaH_2PO_4$, 26 $NaHCO_3$ and 25 glucose, bubbled with 95% $O_2$ and 5% $CO_2$ (pH 7.4). All experiments were performed in transverse slices to avoid the disruption of normal calcium dynamics that is observed close to the stimulation electrode (Bouvier et al., 2016). Slices were visualized using a 40X water-immersion objective (0.75 NA, Axioskop, Carl Zeiss) and infrared optics (illumination filter 750+/- 50nm; CoolSnap Photometrics, Roper Scientific). The recording chamber was continuously perfused at a rate of 4-6 mL/min with a solution containing (mM): 125 NaCl, 2.5 KCl, 1.5$CaCl_2$, 1.8$MgCl_2$, 1.25 $NaH_2PO_4$, 26 $NaHCO_3$, 25 glucose and 10 tricine (a $Zn^{2+}$ buffer, Paoletti et al., 1997) bubbled with 95% $O_2$ and 5% $CO_2$ (pH 7.4). Patch pipettes had resistances between 2.0–4.0 MΩ with the internal solutions given below. Unless otherwise stated, cells were voltage-clamped at -70 mV in the whole-cell configuration. Series resistance was held between 4 and 10 MΩ and compensated with settings of 90% in a Multiclamp 700B amplifier. Whole-cell recordings were filtered at 1-3 kHz and digitized at 10 kHz. Experiments were performed at 32°C (Single Channel Heater Controller, Warner Instruments). 10-20 μM bicuculline methochloride was added to the bath to block GABA-A mediated fast inhibitory transmission. The internal solution contained (in mM): 120 K-Gluconate, 0.5 $K_3$Citrate, 0.5 L (–) Malic acid, 0.008 Oxaloacetic acid, 0.18 α-Ketoglutaric acid, 0.2 Pyridoxal 5-phosphate, 5 L-Alanine, 0.15 Pyruvic acid, 15 L- Glutamine, 4 L-Asparagine, 1 L-Glutathione reduced, 0.5 NAD+, 5 Phosphocreatine $K_2$, 10 Hepes, 0.1 $K_3$EGTA, 4 KCl, 2.2 $K_2HPO_4$, 3.5 NaAcetate, 0.05$CaCl_2$, 2.1 Mg-ATP, 0.4 Na-GTP, 1.4 Na-ATP, pH adjusted to 7.3 with KOH. EPSCs were evoked by stimulating PFs extracellularly by means of a glass pipette (tip diameter 8-12 μm) filled with Hepes-buffered saline. The stimulation electrode was placed at the surface of the molecular layer at 100-500 μm from the recorded PCs. Images were taken every 5 min, experiments showing significant slice movement were discarded. Biphasic stimulation intensity was fixed at the beginning of the experiment (between 1 and 15V; 50 to 100 μs) and remained unchanged during the experiment. Test stimulation was applied at 0.05 Hz. Stimulation consisted of two pulses separated by 50ms, allowing the quantification of PPF. Recordings were made in the vermis of lobules three to eight of the cerebellar cortex. Unless otherwise indicated, long-term potentiation was induced by stimulating the PF beam with 5 pulses at 200 Hz every second for 5 min. During induction PCs were held in current clamp at -70 mV. Long-term depression was induced by stimulating PFs (two pulses at 200 Hz) followed by CF stimulation (four pulses at 400 Hz) 100 ms later, every second for 5 min (Ly et al., 2013). Synaptic plasticity was quantified as the ratio between EPSC charge after induction (mean of 15 sweeps between 30 and 35 min after induction) and control EPSC charge (mean of 15 sweeps immediately before induction). pClamp 9 software (Molecular Devices) was used for data acquisition. Analysis was performed using scripts developed in house with Python 2.6. Statistical significance was determined by using the Wilcoxon rank sum test (R project). All data are shown as mean ± SD.

**Calcium imaging.** All experiments were performed with a custom-built random-access two-photon laser-scanning microscope (Otsu et al., 2008, 2014). In this instrument, both X and Y scanning are operated by acousto-optic deflectors (AODs). These non-mechanical beam-steering devices (A-A Opto-Electronic) can redirect the laser beam in 10 µs. To operate the AODs and run the scanning procedures, a user interface was programmed in LabView (National Instruments). The AOD acoustic frequency drive was generated by a Direct Digital Synthesizer and a fast (10 ns) power amplifier (A-A Optoelectronics). Fast scanning was achieved by generating linear acoustic frequency chirps in both AODs. The spherical lensing effect was automatically compensated by piezoelectric adjustment of the focus. Tunable scanning speed could be achieved up to 0.1 µs per pixel with adjustable regions of interest. Two-photon excitation was produced by an infrared Ti-Sa pulsed Chameleon xr laser (Coherent) tuned to 920 nm coupled to the microscope (Slicescope, Scientifica). The microscope was equipped with a 25x LUMPlanFL/IR objective with 0.95 numerical aperture (Leica microsystems). Fluorescence photons were detected by cooled AsGaP H10769PA-40 photomultipliers (Hamamatsu) in the transfluorescence and epifluorescence pathways, with 641-75 bandpass filter (Omega) for the red morphological channel and a 510-84 bandpass filter (Semrock) for calcium signals. Dwell times of 20-40 µs per point were used. To perform stable long-duration acquisition, registration of the field of view was implemented online between each episode of stimulation by 3D correlation between the image of a small field of view and a previously acquired 3D stack of the same region. This technique allows for the collection of photons from the same point from throughout the recording. Imaging was done in the continuous presence of NBQX (5 µM), bicuculline methochloride (10 µM) (or gabazine (5 µM)) and AM-251 (1 µM). APV (150 µM) and $Zn^{2+}$ (300 nM) were applied for 20 minutes. Experiments were performed at 32°C (Single Channel Heater Controller, Warner Instruments). PF varicosities were identified using a median filter on the peristimulus averaged images. We set a threshold on the area of the objects. After burst stimulations (25 pulses every 15 seconds), the smoothed value of the fluorescence peak was calculated as ΔF/F (ΔF/F = (F-F0) / F0). Only varicosities with relatively large (ΔF > 3 x std(baseline)) and stable signals (ΔF/F < 5% drift between control and drug washout values) were retained for analysis. The DF/F was normalized by the 10 minutes of baseline. Distribution of the baseline, APV, and washout values were taken from the last 5 minutes of each condition. Note that we used 25-30 stimulations of parallel fibers at high frequency as the signal arising from presynaptic NMDAR activation is expected to be very small. This is indicated by the lack any detectable change in transmitter release. In order to maximize the chance to observe such a small signal and faithfully quantify an NMDARs dependent calcium entry in PFs, we perform a longer PF burst than the one used during synaptic plasticity induction, as the goal of this experiment is to test the existence of functional presynaptic NMDARs.

**Compensatory eye movement adaptation.** Male L7cre::GluN1$^{fl/fl}$ and α6cre::GluN1$^{fl/fl}$ mice aged between 10 and 30 weeks and age-matched controls were used in this study. The mice were kept on a 12-hour light/12-hour dark cycle, with ad libitum access to food and water. Behavioral experiments were performed during the light period of the cycle and the experimenter was blind for genotype during experiment and analysis. Mice were prepared for recordings by placing an immobilizing construct (pedestal) on their skulls (Zhou et al., 2015). In short, mice were anaesthetized using isoflurane (initiation 4%, maintenance 2%, with $O_2$), the skin covering the frontal, parietal and interparietal bones was shaved and opened along the rostro-caudal midline. Using Optibond (Kerr, Salerno, Italy) and Charisma (Heraeus Kulzer, Hesse, Germany), a U-shaped holder (brass, 6 × 4 mm) with a magnet inside (neodymium, 4 × 4 mm, MTG, Weilbach, Germany) was fixed on the skull, overlying the frontal and parietal bones. After recovery of >72 hours, mice were habituated to the experimental setup by head-fixing them in a custom-made restraining tube with head holder. Compensatory eye movements were recorded as described before (Schonewille et al., 2010). At least 5 days after surgery, mice were again head-fixed and placed in a restrainer that was positioned in the middle of a turntable surrounded by a cylindrical screen (with a diameter of 60 and 63 cm, respectively). Baseline optokinetic reflex (OKR) and vestibulo-ocular reflex (VOR) were evoked by rotating the screen in the light and table in the dark, respectively (5° amplitude, 0.1–1.0 Hz frequency). The visually enhanced VOR (VVOR), combining visual and vestibular input, was tested by VOR stimulation in the light in a similar manner. Cerebellar motor learning was evaluated in a multiple day paradigm aimed at adapting the gain and phase of the VOR. For four days mice were subjected each day to five 10-minute training sessions of sinusoidal in-phase screen and table rotations at 0.6 Hz, starting on day 1 with both at 5° amplitude, aimed at decreasing the gain of the VOR, followed by in-phase table and screen rotations with screen amplitudes varying from 7.5° (day 2) to 10° (days 3 and 4) to reverse the phase of the VOR. Mice were kept in the dark between training days to prevent active extinction. Before, in between and after the 10 min training sessions VOR is tested (in the dark) to evaluate training effect. Eye movements were recorded at 120 Hz using a CCD camera fixed to the table. Pupil position was obtained using an eye-tracking system (ISCAN) by subtracting the position of a camera-fixed light from that of the center of the pupil. Video calibrations and subsequent eye movement computations were performed with custom-made MATLAB (MathWorks) routines, as described previously (Stahl et al., 2006; Vision Research). To characterize compensatory eye movements, phase and gain were calculated by fitting a sine function to the averaged eye and stimulus velocity traces. Gain was calculated as the ratio of eye-to-stimulus velocity traces. Phase was computed as the difference in degrees between the eye and the stimulus velocity. Consolidation was calculated as the percentage of gain change maintained on day 2 relative to the change achieved during the first day, i.e. $100\% * \Delta gain(day1_{max}-day2_{max}) / \Delta gain(day1_{max}-day1_{min})$. Student's 2-tailed ***t*** tests were used for group comparisons of a single variable (e.g., consolidation). A repeated-measures ANOVA with genotype as the between-subjects factor and frequency or time as the within-subjects factor was used to assess group differences in the eye movement recordings. For all analyses, a *P*-value of less than 0.05 was considered significant.

**Chemicals**. Tricine (10 mM) was used for $Zn^{2+}$ buffering. To obtain 300 nM free $Zn^{2+}$, 60 µM total $ZnCl_2$ was added to 10 mM tricine as described by (Paoletti et al., 1997). All drugs were continuously perfused. Bicuculline methochloride, AM251, and L-NAME were purchased from Tocris Cookson. D-APV was from Ascent Scientific. All other chemicals were from Sigma.

**SI Figures:**

**Figure S1: LTP induction is frequency-dependent and depends on NMDARs containing GluN2A subunits in transverse slices.**

**(A)** Schematic of the Purkinje cell orientation according to the slicing angle. Note that the distance between the surface of the slice and the dendritic arbor of the PC is quite variable. **(B-D)** Transverse slices experimental condition using 5 stimulations at 200 every second, 300 repetitions to induce LTP. **(B)** Representative recordings (averages of 15 sweeps) before (grey) and after (dashed line) LTP induction from (C, D): control (black circle, n = 12 cells), APV (red circle, n = 7 cells), 60ms (grey circle, n = 6 cells), global GluN2A ko (white, n = 7 cells) and his wt littermate (turquoise, n = 6 cells). Note that the induction protocol consists of a burst of 5 PFs stimulations (200 Hz in control condition vs 16.7 Hz using 60ms interval) every second for 5 minutes. **(C)** top: Time course of the normalized EPSC Charge in control, APV, and 60ms. bottom: Time course of the normalized PPR, same condition as in top. **(D)** top: Normalized EPSC charge comparing conditions in (C) and of the GluN2A ko, at t=30-35 min after LTP induction. bottom: Normalized PPR, same condition as in top. **(E-G)** Sagittal slices experimental condition using a burst of 7 PFs stimulations (100 Hz) every second, 300 repetitions (circles), while using 5 stimulations at 200 Hz every second, 300 repetitions to induce LTP (squares). **(E)** Representative recordings (averages of 15 sweeps) before (grey) and after (dashed line) LTP induction from (F-G): control (black circle and grey square), and APV (red circle and purple square). **(F)** top: Time course of the normalized EPSC charge in control conditions (black, n = 9 cells) or in the presence of APV (red, n = 6 cells) using (Piochon et al., 2010) induction protocol (7PFs at 100Hz, 300 repetitions). bottom: Time course of the normalized PPR, same condition as in top. **(G)** top: Normalized EPSC charge comparing conditions in (F) and 5PFs at 200Hz protocol in control (grey square, n = 7 cells) and APV (purple square, n = 6 cells), at t=30-35 min after LTP induction. bottom: Normalized PPR, same condition as in top. PPR was not changed after synaptic plasticity induction for any of those conditions. Statistical significance was tested using Wilcoxon test.

**Figure S2: High frequency stimulation of parallel fibers induces NMDAR-dependent nitric-oxide synthesis.**

**(A)** Representative paired pulse EPSCs before (grey) and after (dashed line) LTP induction in the control condition (black circle) and during application of 100 µM L-NAME (blue circle). LTP was induced using 5 stimulations at 200 every second, 300 repetitions **(B)** top: Time course of the normalized EPSC charge in control conditions (black, n = 12 cells) and in presence of L-NAME (blue circle, n = 7 cells). bottom: Time course of the normalized PPR, same condition as in top. **(C)** top: Normalized EPSC charge comparing conditions in (B) at t=30-35 min after induction. bottom: Normalized PPR, same condition as in top. PPR was not changed after synaptic plasticity induction for any of those conditions.

**(D)** Top panel: Schematic representing the recording of NO signals by using electrochemical detection after burst stimulation of parallel fibers (see Methods). Time course of the absolute current in control conditions (black, n = 20 slices), in presence of L-NAME (blue, n = 12 slices), in APV (red, n = 15 slices), and using GC-GluN1 ko (green, n = 14 slices). **(E)** Box plot comparing each condition in (D) the last minute of the induction (t=240-300 s after the beginning of burst stimulations, 25-30 PFs stimulations at 200Hz, every second for 5 minutes) and the last minute of the baseline. Statistical significance was tested using Wilcoxon test.

**Figure S3. Cerebellum-dependent compensatory eye movements after genetic ablation of NMDARs.**

**(A)** Schematic representation of the behavioral experiment to determine the characteristics of the compensatory eye movements by measuring optokinetic reflex (OKR), vestibulo-ocular reflex (VOR), and visually-enhanced (VVOR). Sinusoidal rotation of a drum covering the entire visual field generated a tracking OKR, rotation of the turntable in the dark resulted in a compensatory VOR, and in the light the VVOR. **(B)** Mice with Purkinje cell specific deletion of GluN1 (N = 12 mice) have normal gain and phase values for all forms of compensatory eye movements, except for a difference in the phase of the VOR (N = 12 mice for controls, $p = 0.010$, $\Delta$phase = 1.1 ± 0.1°, all other $p > 0.5$). **(C)** Ablating GluN1 from GCs specifically (n = 13 mice) only affected the phase of the VOR significantly compared to controls (N = 12 mice, $p = 0.043$, $\Delta$phase = 6.2 ± 2.2°). Error bars denote SEM (please note that error bars can fall within symbols), * $p < 0.05$, ** $p < 0.001$. Statistical significance was tested using an ANOVA for repeated measurements.

Fig. S1. Schonewille et al.

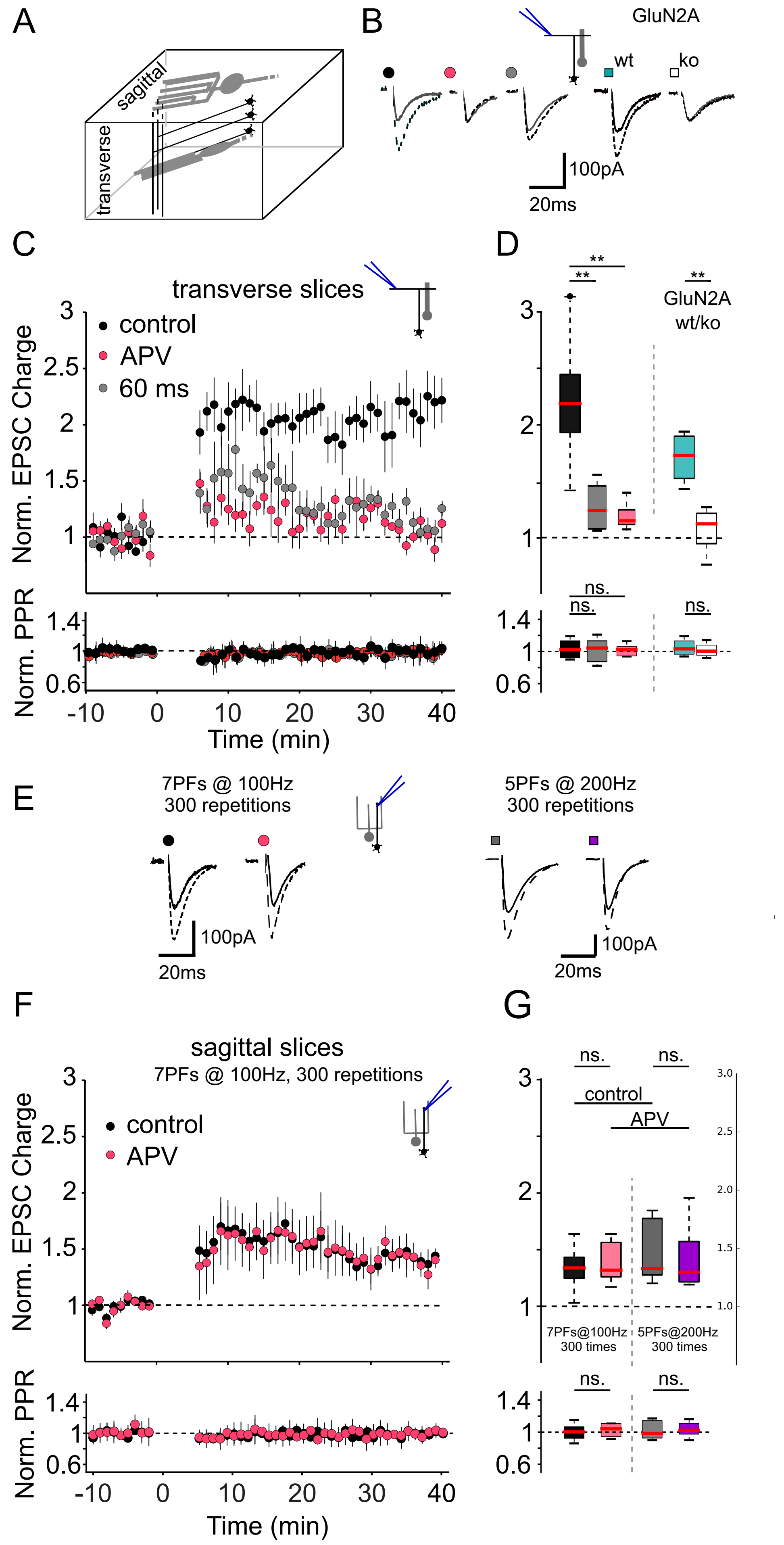

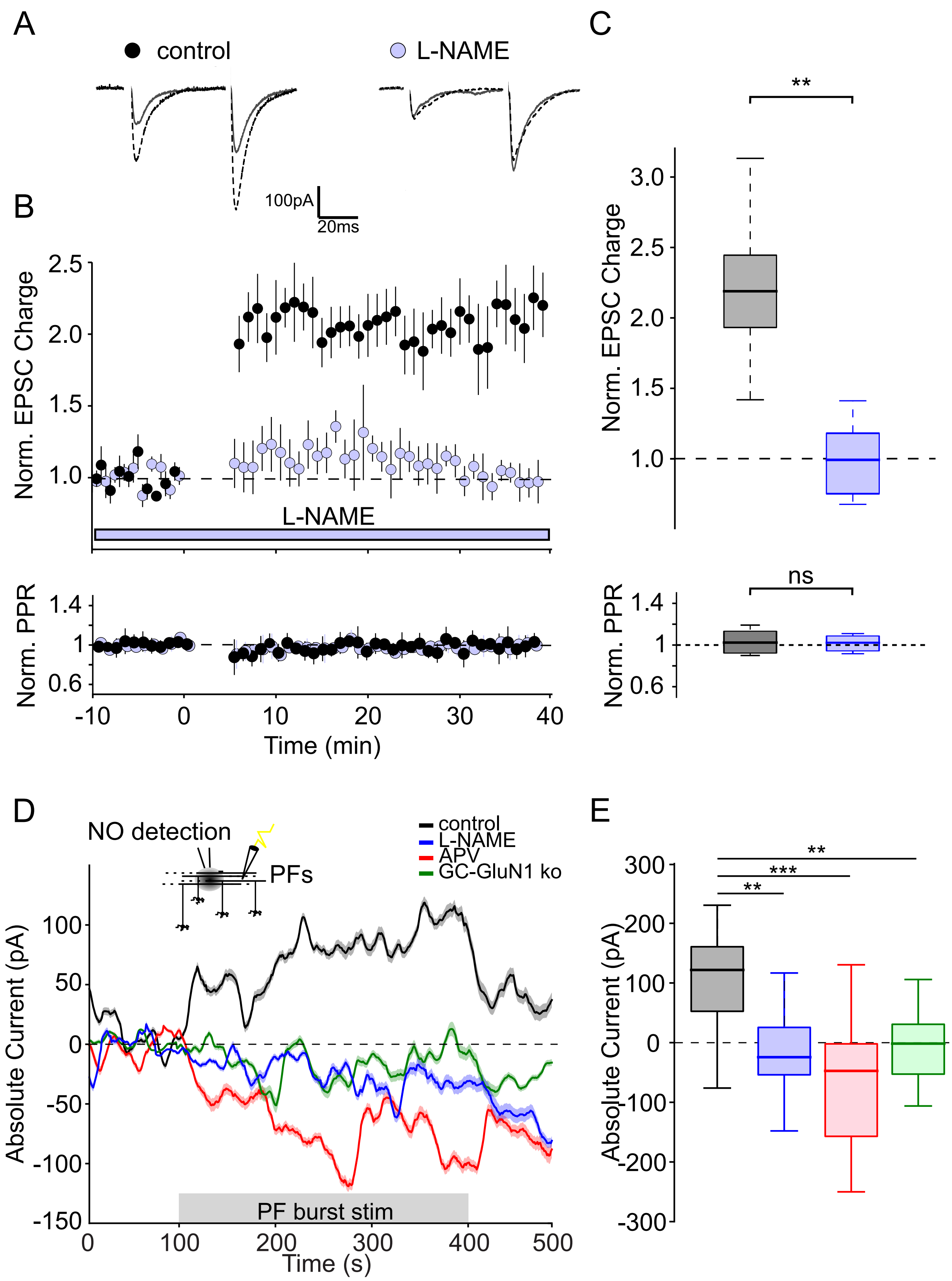
A
control
L-NAME
100pA
20ms
B
Norm. EPSC Charge
2.5
2.0
1.5
1.0
L-NAME
Norm. PPR
1.4
1
0.6
-10
0
10
20
30
40
Time (min)
C
**
Norm. EPSC Charge
3.0
2.5
2.0
1.5
1.0
ns
Norm. PPR
1.4
1
0.6
D
NO detection
PFs
control
L-NAME
APV
GC-GluN1 ko
Absolute Current (pA)
100
50
0
-50
-100
-150
PF burst stim
0
100
200
300
400
500
Time (s)
E
**
***
**
Absolute Current (pA)
300
200
100
0
-100
-200
-300

Fig. S3. Schonewille et al.

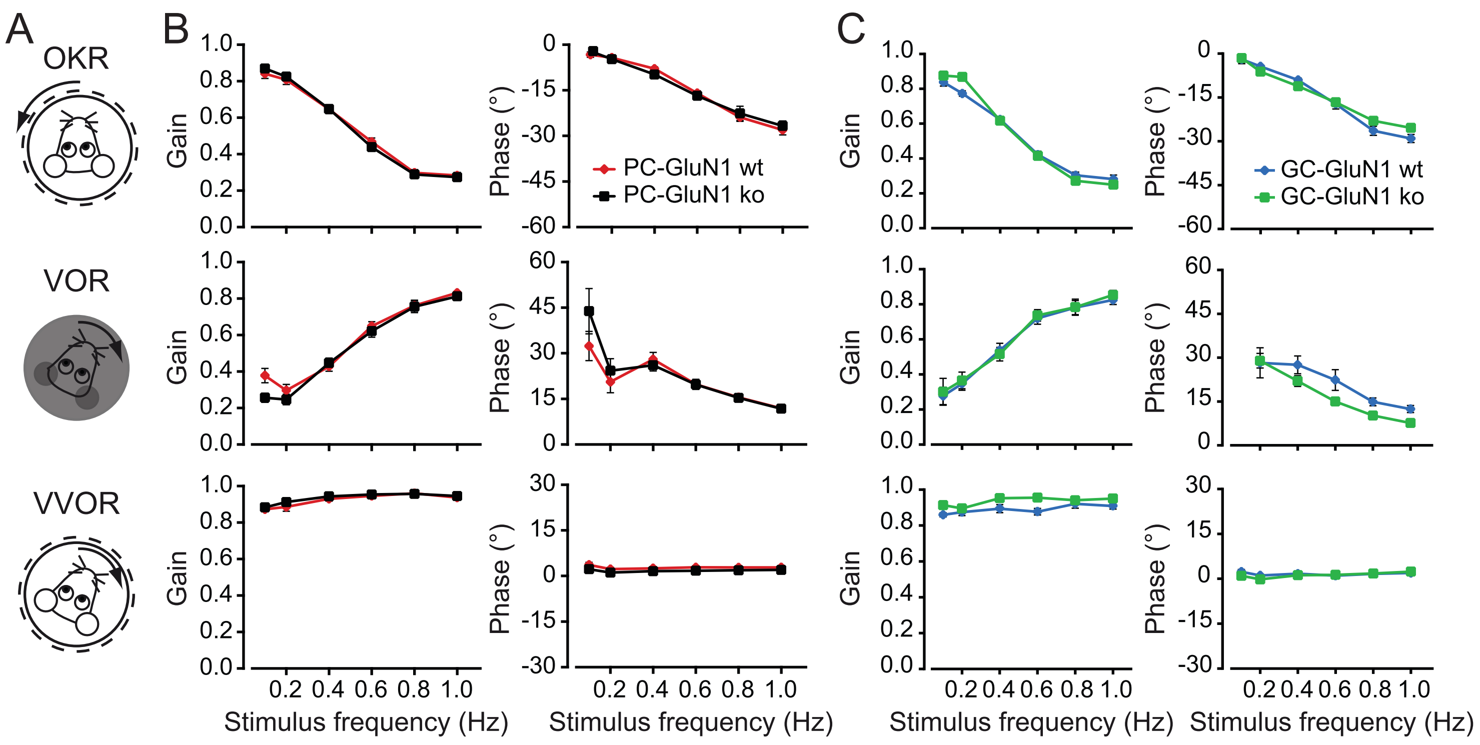
